# The effects of artificial light at night (ALAN) on the gaping activity and feeding of mussels

**DOI:** 10.1101/2023.03.10.532051

**Authors:** Eleni Christoforou, Davide Dominoni, Jan Lindström, Christina Diamantopoulou, Jakub Czyzewski, Nosrat Mirzai, Sofie Spatharis

## Abstract

Artificial light at night (ALAN) is one of the most widespread forms of environmental pollution. Studies on terrestrial organisms have shown that the effects of ALAN can be pervasive, and importantly, can depend on the colour (i.e., wavelength) of light. ALAN also affects marine environments as it is present in more than 22% of the world’s coastlines and can reach depths of up to 100m. However, the impact of different colour ALAN on coastal marine organisms is under-investigated. In this study, we tested the effects of different ALAN colours on *Mytilus edulis*, a widespread coastal bivalve known for its high phytoplankton clearance capacity amongst other valuable ecosystem services. Using a lab-based valvometry system, we recorded the impact of red, green, and white ALAN on gaping activity patterns and phytoplankton clearance capacity of individual mussels and compared these to dark night control. Mussels exhibited a semi-diel activity pattern in both proportion of time open and gaping frequency. Although ALAN did not have significant effects on the proportion of time open it did significantly affect the gaping frequency. This effect was colour-specific with red and white ALAN resulting in lower activity compared to the dark night control but there were no effects on the phytoplankton consumption. Under green light, however, mussels showed a higher gaping frequency and reduced phytoplankton consumption with increasing time spent open compared to the other ALAN treatments and the dark control. Our findings suggest that ALAN does have colour-specific effects on mussels and indicate the importance of further investigating the physiological mechanisms behind these patterns, and their potential ecological consequences.

## 1 Introduction

Due to the expansion of human population and urbanisation, Artificial Light at Night (ALAN) has become a dominant form of environmental pollution, with worldwide lit areas increasing by 2.2% annually (Kyba et al. 2017). This is inevitably affecting living organisms which are evolutionarily adapted to rely on circadian rhythms for essential biochemical, physiological and behavioural processes (Pittendrigh 1960, Naylor 1999, Sharma 2003). For instance, ALAN has been shown to shift morning activity of urbanised birds (Dominoni et al. 2013), disrupt the locomotory activity of sand isopods (Duarte et al. 2019) and decrease the swimming activity of fish inhabiting urban streams (Miner et al. 2021). Impacts induced by alterations to these natural processes may scale up from individual to community level (Maggi and Benedetti-Cecchi 2018, Garratt et al. 2019, Hölker et al. 2021) and have negative effects on the functioning of ecosystems (Zapata et al. 2018). Research on this topic is rapidly expanding, but it is currently largely focused on terrestrial and freshwater ecosystems (Bedrosian et al. 2013, Brüning et al. 2018, Tidau et al. 2021). Yet, it has been shown that about 1.9 million km^2^ of shallow (up to 1m) coastal areas are affected by biologically important ALAN, which can reach depths of up to 100m (Davies and Smyth 2018). The communities of marine organisms serve important environmental functions by providing invaluable ecosystem services such as regulation of water quality and food web stability (Prather et al. 2013). Therefore, more focus should be directed on coastal marine ecosystems where ALAN can be impactful to aquatic communities (Davies et al. 2020, Smyth et al. 2021) and where studies are currently limited.

Bivalves are vital for the healthy functioning of coastal ecosystems (Prather et al. 2013, van der Schatte Olivier et al. 2018) as they are ecosystem engineers (Prather et al. 2013). Through biofiltration and biodeposition, they provide invaluable ecosystem services by facilitating the nutrient cycle (Prather et al. 2013, Kent et al. 2017), controlling phytoplankton abundance and composition, and thereby remediating the impacts of eutrophication and algal blooms (Prins et al. 1998, Tantanasarit et al. 2013, van der Schatte Olivier et al. 2018). Generally, bivalves show higher nocturnal than diurnal activity, represented by increased valve movement, greater valve gape angles, higher exhalant pumping (Robson et al. 2010a), longer duration of open gape (Kobak and Nowacki 2007, Gnyubkin 2010), higher filtration (Hills et al. 2020) and growth rates (Strömgren 1976, Nielsen and Strömgren 1985). As the regulation of diel cycles in these processes is strongly regulated by light, we could expect ALAN to disrupt them, possibly also affecting the associated ecosystem services provided. It is therefore concerning that coastal sessile invertebrates, like mussels, can be especially vulnerable to light pollution due to direct and continuous exposure (Bolton et al. 2017, Gaston et al. 2017) to urban settlement, coastal street and harbour lighting (Zissis and Bertoldi 2018). However, studies on the effects of ALAN on bivalves are currently lacking.

Importantly, it has been suggested that because organisms are generally sensitive to specific wavelengths of light, altering the colour of light installations might minimise the impact of ALAN on wild species. Recently, the development of light-emitting diode (LED) technology (Cho et al. 2017, Zissis and Bertoldi 2018) has provided an energy-efficient and affordable opportunity to shift to ALAN wavelengths that are less stressful to wildlife (Gaston et al. 2017). For instance, red is suggested to be used near organisms with sensitivity or attraction to short wavelengths like blue and green (Spoelstra et al. 2015). Such organisms include bats (Spoelstra et al. 2015), sea turtles (Miller and Bretschneider 2006) and corals (Ayalon et al. 2019). Research on migratory seabirds and mice recommend the use of green light to minimise the effects of ALAN (Poot et al. 2008). This variation in the proposed light spectra is attributed to the organisms’ diverse sensitivity to light wavelengths due to differences in the evolutionary development of their photoreceptive cells and organs (Von Salvini-Plawen 2008, Alaasam et al. 2021). As consensus is yet to be met on the most appropriate ALAN wavelengths for the conservation of coastal wild populations, more research is necessary on dominant coastal organisms such as bivalves that are major contributors to coastal ecosystem services.

Epifaunal bivalves possess photoreceptor organs on their mantle or even light-sensitive multi-cellular eyes (Morton 2008, Von Salvini-Plawen 2008, Audino et al. 2020). Their main role is to induce an instant anti-predatory response when a change is detected in the light intensity of the organisms’ immediate environment (Wilkens 2008). These photoreceptors are also involved in the regulation of circadian rhythms displayed by bivalves (Ortmann and Grieshaber 2003, Garci et al. 2008, Gnyubkin 2010) however information on bivalves’ spectral photosensitivity is scarce and mostly centred around scallop species, which have the most complex eyes, and show peak absorbance at 480-504nm and 513-549nm (Cronly-Dillon 1966, Speiser et al. 2011). Scallops species inhabiting deeper waters are more sensitive to longer wavelengths in comparison to coastal scallops (Speiser et al. 2011). Hence, the range in sensitivity seen in different species could be attributed to the light attenuation and wavelength absorption at different depths and water conditions. Comparative studies on the effects of different wavelengths on mussels point to an overall avoidance of coloured light and preference to dark areas (Kobak and Nowacki 2007). A negative effect of red fluorescent light was expressed as lower growth rate (Nielsen and Strömgren 1985) and greater sensitivity of the photoreceptor cell responsible for the shadow reflex (Cornwall and Gorman 1983); however, research on the effect of ALAN and different ALAN wavelengths on the gaping activity and feeding of bivalves is lacking. It is critical to investigate these responses as they are directly linked to the important ecosystem functions provided by bivalves such as the control of phytoplankton abundance and nutrient cycling.

The aim of this study was to determine whether the gaping activity and phytoplankton consumption of mussels can be impacted by ALAN, and whether these effects are wavelength-specific. To achieve this, we performed a laboratory experiment exposing blue mussels, *Mytilus edulis* to wavelengths relevant for coastal illumination and conservation, namely green, red, and white ALAN, compared to a dark night control. The responses studied included the circadian rhythm of gaping activity (i.e., proportion of time mussels were open and the gaping frequency) and the phytoplankton consumption by mussels. We expect that in the presence of ALAN, mussels will be less active and filter less phytoplankton compared to the dark control. We hypothesise that mussels might be more sensitive to white ALAN as they are adapted to sense changes to daylight. The relationship between gaping activity and phytoplankton clearance capacity, and how this depends on the ALAN colour was also investigated. We would not anticipate any changes to this relationship as we would expect both responses to change in the same manner within each colour treatment.

## 2 Methods

### 2.1 Experimental design

The effect of different ALAN wavelengths (green, red, and white) on (a) the circadian rhythm of the gaping activity of mussels including the proportion of time open and gaping frequency (number of times a valve opened/closed) and (b) the phytoplankton consumption by mussels were investigated experimentally. For the monitoring of gaping activity, mussels were continuously fed with a peristaltic pump for 14 consecutive days and their proportion of time open and gaping frequency were measured every minute through the whole period using a custom-made valvometry system. The phytoplankton consumption, over 24 hours, was measured on the first and last day of the experiment when the mussels were fed only once in the first hour. To account for possible variation due to the reproductive cycle (Fernández et al. 2015), the experiment was run twice for each of three reproductive stage (pre, during and post spawning). Each run consisted of 12 mussels for a total sample size of 72 individual mussels (3 mussels per treatment x 4 treatments x 2 runs per reproductive stage x 3 reproductive stages).

### 2.2 Collection and acclimation

Rope grown *Mytilus edulis* mussels (6.28cm ± 0.34SD, n=72) were collected by Fassfern Mussels staff from Loch Eil, Scotland (56°51**’**14**’’**N,5°11**’**33**’’**W) in 2020 and provided to us within the same day. Mussel energy storage, condition index and weight is affected by food availability and reproductive stage which depend on seasonality (Okumuş and Stirling 1998, Fernández et al. 2015). To account for this variability in the mussels’ physiological state, they were collected in winter (January 22^nd^, SST: 7°C, salinity: 23psu), summer (August 23^rd^, SST: 14°C, salinity: 25psu) and autumn (October 4^th^, SST: 12°C, salinity: 23psu), which are representative of the pre, during and post spawning stages, respectively (Fernández et al. 2015).

In the laboratory, epibionts such as barnacles, macroalgae and epiphytic diatoms were scraped off the mussel shells to reduce interference with gaping activity and feeding. Mussels were then placed in 5L artificial salt water (32ppm) aquariums, located in the light-sealed boxes under 12hour day: 12hour night photoperiod, and were starved for 48 hours to standardise the mussels’ hunger level prior to the treatments. After depuration, the valvometry system was attached to 12 experimental mussels across the four ALAN treatments which were individually placed in experimental cylindrical glass vessels (height: 25cm, diameter: 7.3cm) filled with 550ml of artificial salt water (32ppm). They were then allowed to acclimate for the first 24 hours. Experimental vessels, containing the mussels, were constantly aerated, and temperature was maintained between 12-14°C by submerging the experimental vessels in water baths that were fed by a common thermoregulated water tank. *Tetraselmis* sp. microalgal monoculture (S1) was provided through a peristaltic pump for 15 mins every three hours, for a total phytoplankton concentration of 3×10^6^cells/L. Mussels were then starved for another 24 hours before the initiation of the experiment.

### 2.3 Light exposure

The experiment involved four light treatments (dark control and red, green, and white ALAN), each performed in a light-sealed box under 12-hour day: 12-hour night photoperiod. In each treatment, a white LED light (496-627nm with a peak at 548nm) was used to simulate day light. For the night treatments we used green (505-586nm, peak at 536nm), red (510-664nm, peak at 634nm) and warm white (503-620nm, peak at 558nm) LED as ALAN sources (see Supplementary material S2 for the spectrum profiles). A dark night (0 lux) treatment box served as the control. The day-time lux levels were standardised across all treatment boxes at an intensity of 3702.5lux ± 249.2SD according to the lux range of a cloudy day (1000-10,000 lux). The ALAN lux levels were standardised at 19.86lux ± 0.5 SD across the ALAN treatment boxes to approximate the illuminance of the average street ALAN (15 lux) (Gaston et al. 2013) and the lux levels measured at the water surface of a coastal environment (5-21.6 lux) (Davies et al. 2015). The light sources were positioned 40cm above the water surface and their illuminance was measured with a LI-210R photometric sensor (LI-COR, USA) at the water surface.

### 2.4 Recording of gaping activity

To investigate the effect of ALAN wavelengths on the circadian rhythm in gaping activity of mussels, acclimated mussels were placed individually in three experimental vessels in each of four light-sealed treatment boxes. The proportion of time open and gaping frequency were tracked and recorded every minute (Robson et al. 2009) for the duration of the experiment. This was achieved by a custom-made valvometry system (S3), developed by the Bioelectronics Unit of the School of Biodiversity, One Health & Veterinary Medicine at the University of Glasgow, according to previous bivalve studies (Andrade et al. 2016, Comeau et al. 2018, Clements and Comeau 2019). Mussel valves were recorded as closed when they were in contact at the posterior end, otherwise they were recorded as open (Figure 1).

**Figure 1:**
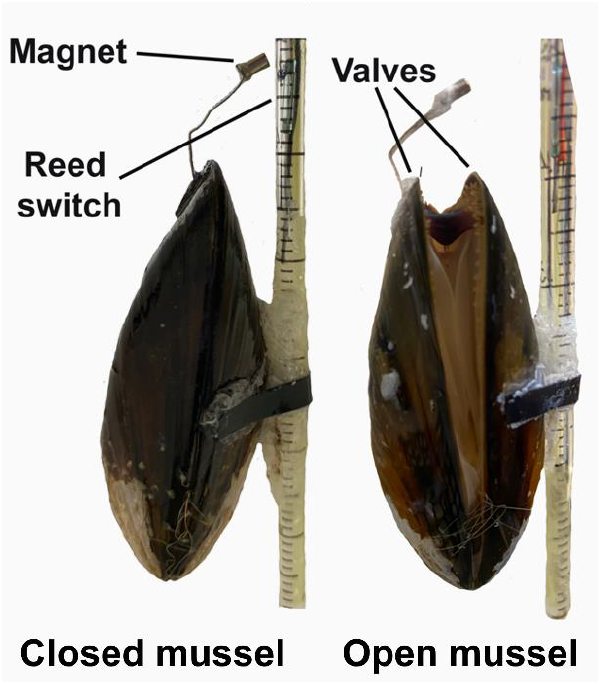
Photos of mussels equipped with the valvometry system where the magnet, the reed switch and the valves of the mussels are labelled. The mussel on the left represents a closed mussel where the magnet would trigger the reed switch, close the circuit and a signal would be recorded while the mussel on the right would be open and not trigger the reed switch. For more details regarding the valvometry system see Supplementary information S3.

It has been suggested that mussel activity and feeding could be affected by the time of feeding in laboratory experiments (Robson et al. 2010a). To eliminate any such effects, we fed *Tetraselmis* sp. to the mussels in a semi-continuous manner for 15 mins every three hours using a peristaltic pump. This added up to 982ml of artificial saltwater, after 48 hours, with a total phytoplankton concentration of 3×10^6^ cells/L. At the end of each 48-hour interval, within the last hour of the daytime period (17:00-18:00), the gaping status recording was paused to allow for the renewal of the artificial salt water. During that time, the experimental vessels were rotated within their treatment boxes to allow for a more homogeneous light exposure.

To assess whether ALAN affected circadian rhythms of gaping activity, we first calculated the proportion of time open, using the following equation:

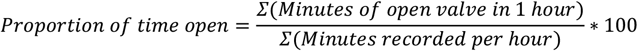

The sum of the minutes that the valve was open per hour was divided by the minutes of available recording per hour to account for any incomplete hours which occasionally occurred due to system errors. The value is multiplied by 100 to express the proportion of time open as a percentage.

We also calculated the gaping frequency using the following equation:

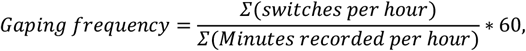

where switches are the transitions from close to open valve and vice versa per hour. These are divided by the number of minutes recorded and then multiplied by 60 for a standardized count within an hour.

### 2.5 Phytoplankton consumption

Phytoplankton consumption was measured on days 1 and day 14 of the experiment. For the consumption quantification, 800ml of artificial saltwater containing 3×10^6^cells/L (T_0h_) *Tetraselmis* sp. monoculture was added to the experimental vessels, each containing a single mussel. After 24 hours, a 50ml water sample was collected, to measure the final concentration (T_24h_) according to Christoforou et. al. (2020). In brief, samples were preserved in amber glass bottles with lugol iodine solution and filtered using SartoriusTM Cellulose Nitrate Membrane Filters. The filters were then dried, made transparent using immersion oil and the cells were visualised and counted using a light microscope camera at x40 magnification. Phytoplankton consumption was calculated as the percentage of phytoplankton consumed by each individual mussel, using the following equation:

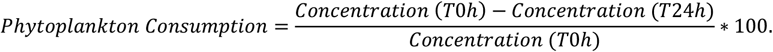

### 2.6 Data analysis

To determine the effect of ALAN treatments on circadian patterns in mussels’ gaping activity, we fitted two General Additive Mixed Models (GAMMs) using the R mgcv package (v1.8-33) (Wood 2011). The two response variables for the two models were proportion of time open and gaping frequency and the explanatory variables were the hour of the day which was modelled as a smooth term, the ALAN treatment (categorical factor variable with four levels: green, red, white, and dark) interacting with period (categorical factor variable with two levels: day and night), the reproductive stage (pre, during and post spawning), the covariate experimental day, and the occurrence of water exchange on the day (factor yes/no). The proportion of time open ranged between 0 and 1 and was modelled as beta distribution, which is most suitable for proportional data, after transforming the values with the following formula to allow the use of values which are exactly zero or one:

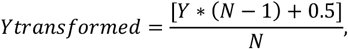

where Y is proportional value (i.e., the proportion of time open),, *Ytransformed* is the transformed value (i.e., the transformed proportion of time open), and N is the sample size (Smithson and Verkuilen 2006). The zero-inflated count data of gaping frequency were modelled using a Poisson error distribution as this had the highest deviance explained (22%) compared to other tested distributions (quasi-Poisson, negative binomial, and zero-inflated Poisson).

Phytoplankton consumption comprised of proportional data and was transformed using the aforementioned transformation equation where Y would be the proportion of phytoplankton consumed. We modelled phytoplankton consumption using a generalized linear mixed model (GLMM) with beta distribution. The explanatory variables included the proportion of time open and gaping frequency interacting with the ALAN treatments, the reproduction stage, the experimental day, the acclimation period, and the initial phytoplankton concentration. The model was fitted using the glmmTMB package (v1.0.2.1) (Brooks et al. 2017). Outliers detected in the phytoplankton consumption data were omitted from the analysis (see Supplementary material S4 for details on outlier removal procedure).

In all models, the mussel ID nested within the run number was used as a random effect to account for individual mussel variation within the runs. The analysis was conducted in R v.4.0.4 (R Core Team 2021) and model selection was performed using model AIC comparisons. To pinpoint significant differences between the treatments at the picks and troughs seen during the activity cycles, we performed pairwise comparisons tests using the Tukey method in the emmeans package v1.5.2-1 (Lenth 2018). This was achieved by setting the hour variable as a factor and running the selected simpler model as a GLMM.

## 3. Results

### 3.1. Effects of ALAN on the valve gaping activity of mussels

The circadian activity in the proportion of time open showed a clear semi-diel pattern (Figure 2A) with hour of the day having a significant effect across all ALAN treatments (Table 1, p<0.001 see model output (A) in Supplementary material (S5)). Specifically, the proportion of time open peaked at early daytime (7:00 hrs) and then in early night-time (19:00 hrs) with the night peak being the highest (Figure 2A). There was no significant difference between the ALAN treatments nor against the dark control.

**Figure 2:**
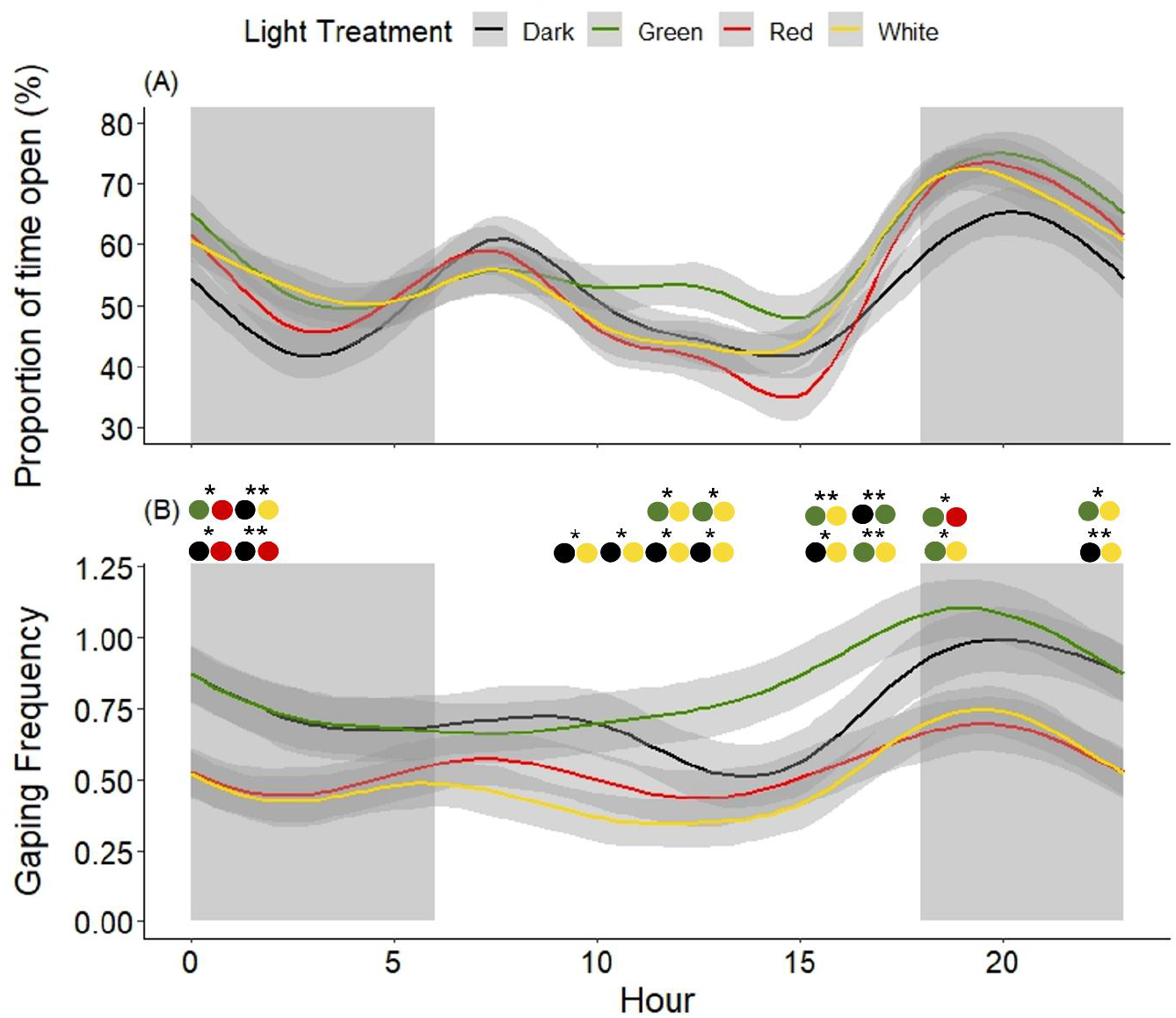
The mussel gaping activity expressed as (A) the proportion of time open and (B) gaping frequency across the ALAN treatments (dark, green, red, and white). Each line is plotted through a 24-hour period and represents the mean of all experimental days with the 95% confidence interval (grey bands). Day time (white area) was from 6:00-18:00 and night-time (grey area) from 18:00-6:00. The asterisks above panel B indicate which pairs of treatments (colour dots) were significantly different from each other at that time of the 24-hr cycle (p<0.01** and p<0.05*).

**Table 1:** The explanatory variables which, after AIC model selection, were found to explain significant variation in the mussel (A) proportion of time open, (B) gaping frequency and (C) phytoplankton consumption. The ΔAIC indicated the difference in the AIC when the explanatory variable was dropped from the best-supported model. The selected model outputs can be found in Supplementary material (S5).

| **Response (bold)** and explanatory variables (not bold) | **\|Δ AIC\|** |
| --- | --- |
| **(A) Proportion of time open** |  |
| Experimental day | 271.8 |
| Water change | 216.3 |
| s (Hour, by=ALAN treatment) | 441.6 |
| **(B) Gaping frequency** |  |
| ALAN treatment * day/night | 10.9 |
| Experimental day | 199.9 |
| Water change | 43.1 |
| s (Hour, by=ALAN treatment) | 431.7 |
| **(C) Phytoplankton Consumption** |  |
| Proportion of time open * treatment | 8.4 |

Gaping frequency also peaked at early night-time at 19:00 hrs and showed a similar but less pronounced semi-diel pattern (Figure 2B). ALAN treatment significantly affected the gaping frequency with higher activity under the green ALAN 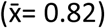 and the dark control 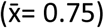 in comparison to the red 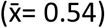 and white ALAN 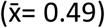. Significant pairwise differences between the treatments at each hour of the experiment are indicated in Figure 2B.

### 3.2. Effects of ALAN on phytoplankton consumption

The proportion of time open significantly affected phytoplankton consumption by the mussels (Figure 4A). This relationship depended on the ALAN treatment (see interaction between ALAN treatment and proportion of time open in Table 1). Specifically, in the dark, red, and white ALAN treatments’ mussel consumption increased by 14.4%, 18.3% and 56.1%, respectively, with increasing proportion of time open whereas the reverse was observed under the green ALAN treatment (−27.0%). There was no significant difference in the overall phytoplankton consumption between the treatments (Figure 4B).

**Figure 4.**
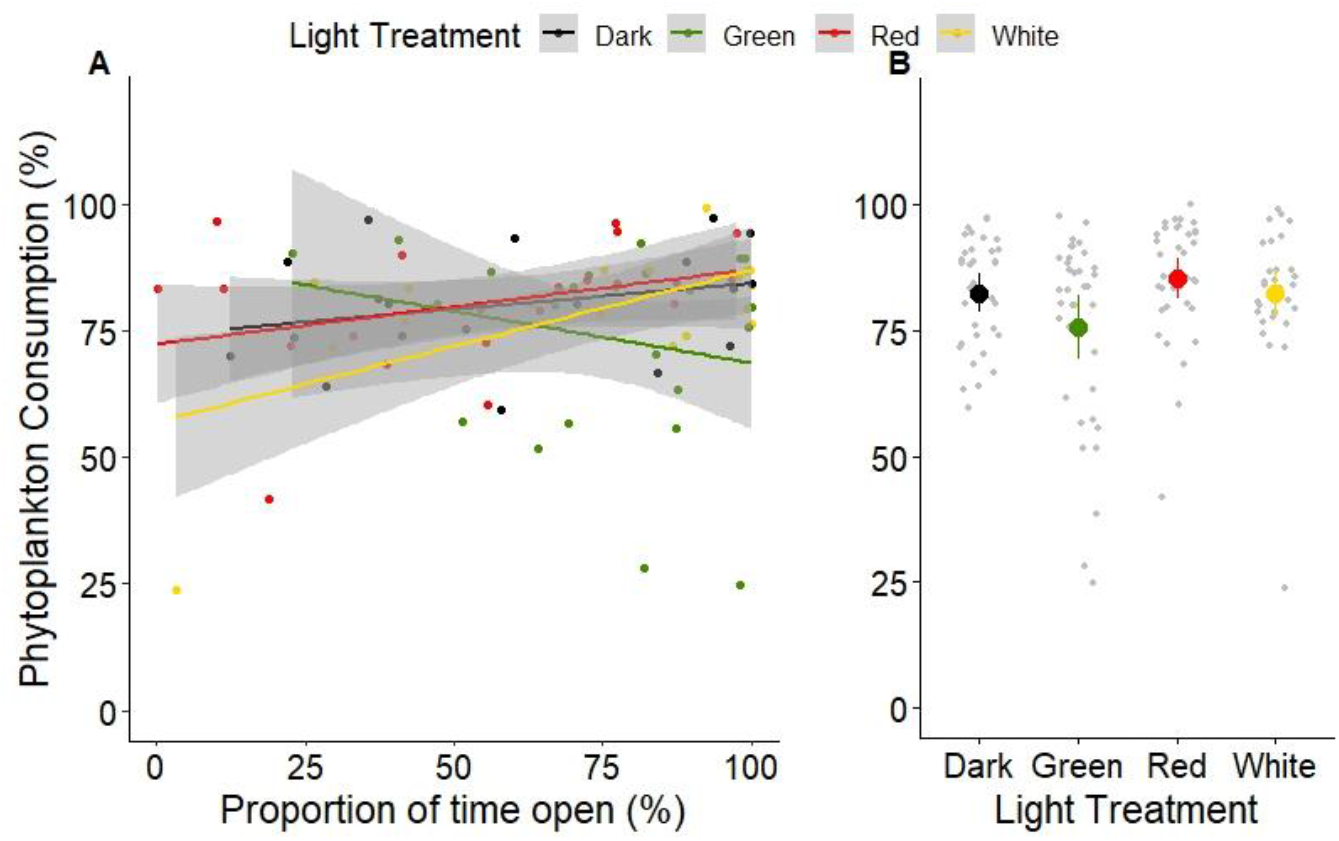
(A) The relationship between mussel phytoplankton consumption and proportion of time open, both expressed in percentages, across the ALAN treatments (dark, green, red, and white) represented by a regression line and the 95% confidence intervals (grey bands). (B) The Mean ± SE of phytoplankton consumption by mussels over 24 hrs, expressed across the ALAN treatments. Experimental day had no significant effect on the clearance capacity of mussels thus data from both experimental days (days 1 and 14) were pooled together.

## 4. Discussion

This is, to our knowledge, the first study investigating the effect of ALAN on bivalves and comparing the impacts of different ALAN colours. We found that the mussel activity expressed both as proportion of time open and gaping frequency follows a semi-diel pattern. Neither the proportion of time open patterns nor the phytoplankton consumption was affected by ALAN. However, mussels exposed to white and red ALAN displayed significantly lower gaping frequency than the dark control by 32.4% and 40.8%, respectively. Those exposed to green ALAN were more active by 10% than the dark control. Interestingly, the relationship between proportion of time open and phytoplankton consumption was positive at the white and red ALAN and the dark control, but it was reversed under green ALAN.

The semi-diel pattern observed in the proportion of time open mirrors the time of tides at the site of collection of mussels (see figures in Supplementary material S6). Tide-related activity patterns have been also observed in fully submerged (i.e., deprived of tidal information) clams (Williams and Pilditch 1997), oysters (Tran et al. 2020), and mussels (Pampapathi 1954, Zaldibar et al. 2004, Gnyubkin 2010). Ortmann and Grieshaber (2003) have suggested that the relationship between tidal activity and the fluctuation of phytoplankton levels in the field could determine the gaping activity of bivalves. Our findings suggest that intertidal *Mytilus edulis* mussels can still follow a circa-tidal activity pattern despite being provided with a constant phytoplankton supply indicating an endogenous behaviour (Tran et al. 2020, Bertolini et al. 2021). This circa-tidal behavioural rhythm observed may be attributed to the anticipation of phytoplankton arrival brought in with the tide (Williams and Pilditch 1997, Riisgård et al. 2006, Saurel et al. 2007).

In agreement with our hypothesis, we found that the gaping frequency was reduced under white ALAN compared to the dark night control. This indicates that mussel activity patterns might indeed be sensitive to even very low white light intensities (lux). We also found that gaping frequency was significantly reduced under red ALAN, and this agreed with a previous study by Nielsen and Strömgren (1985) who found that red light affected coastal mussels by reducing their growth rates, which is usually a sign of suboptimal physiological functioning. Although the reduced gaping frequency of mussels under red and white ALAN was not linked to a reduction in phytoplankton consumption, further investigation is necessary on possible effects on other physiological and behavioural functions, and reproduction.

The increased gaping frequency of mussels under green ALAN is in line with previous studies that have linked high gaping frequency to stress (Andrade et al. 2016). It was suggested that bivalves would open to test the ambient conditions and close at the presence of a disturbance (Curtis et al. 2000, Kobak and Nowacki 2007) which could be the case in the presence of green ALAN. High gaping frequency can be detrimental to the individuals’ physiology and fitness, as increased opening and closing is energetically costly, can affect the heart rate of bivalves (Curtis et al., 2000) and increase chances of predation (Kobak and Nowacki, 2007). Under green light, the mussels that spent more time open consumed less phytoplankton, in contrast to the other light treatments and previous studies that showed positive relationship between the activity and feeding (Jørgensen et al. 1988, Frank et al. 2007). This indicates that mussels under the green treatment used the time spent with the valves open for a different function rather than feeding. Indeed, some studies on mussels have argued that the valve gape status cannot be used as a proxy for the feeding activity (Jørgensen et al. 1988, Frank et al. 2007, Macdonald et al. 2009). Other physiological activities that would require a valve opening include breathing, rejection of faeces and pseudofaeces, foot protrusion for movement and attachement via byssus threads (Gosling 2003, Robson et al. 2010b). It is thus possible that under green ALAN, mussels kept the valve open for e.g., protruding their foot in an attempt to move away from the green ALAN, (even though displacement was impossible due to the attachment to the valvometry system) instead of filtering efficiently for phytoplankton during that time. The attempt to escape from green ALAN is supported by Kobak and Nowacki (2007) who showed that zebra mussels displayed equal aversion to green, blue, red, and white coloured light (daytime exposure, not ALAN), after 24 hours exposure to the same light regime, by moving to a shaded area. This possible explanation merits further investigation, for instance by video recording mussels’ behaviour during ALAN exposure.

Our findings show that mussels exhibit sensitivity to ALAN compared to the dark night. Mussels were less active when exposed to white and red ALAN while mussels exposed to green ALAN show a higher gaping frequency and a negative relationship between proportion of time open and phytoplankton consumption. It can be speculated that increased activity under green light that is not directed towards feeding could entail major energetic costs to the animals in the long term. Additionally, Diamantopoulou et al. (2021) have shown that green and red ALAN can enhance phytoplankton growth and alter its assemblage with opportunistic, bloom-forming diatom species dominance. Such changes in phytoplankton concentrations and composition in conjunction with alterations in the activity and filtration capacity of mussels under green and red ALAN could change the balance maintained by these filter feeding organisms and have noxious effects on the ecosystem, leading to harmful algae bloom formation and subsequent devastating ecological impacts. Because of these potentially important implications, further experimental work is required to establish what the long-term consequences of changes in mussels’ behaviour under ALAN are on mussel physiology, reproduction and growth.

## Acknowledgements

We would like to thank The A.G. Leventis Foundation and Nissad Development Company for their financial support (sponsors had no further involvement in this study). We would like to thank Fassfern Mussels for providing the mussels and Prof. Neil Metcalfe and Dr. Eirini Kaiserli for the provision of some necessary equipment facilitating the implementation of the experimental as well as the IBAHCM aquarium staff for their help in setting up the aquaria used.

## Notes

### Competing Interest Statement

The authors have declared no competing interest.

## References

Alaasam, V. J., M. E. Kernback, C. R. Miller, and S. M. Ferguson. 2021. The Diversity of Photosensitivity and its Implications for Light Pollution. Integrative and Comparative Biology 0:1–12.

Andrade, H., J. C. Massabuau, S. Cochrane, P. Ciret, D. Tran, M. Sow, and L. Camus. 2016. High frequency non-invasive (HFNI) bio-sensors as a potential tool for marine monitoring and assessments. Frontiers in Marine Science 3:1–10.

Audino, J. A., J. M. Serb, and J. E. A. R. Marian. 2020. Hard to get, easy to lose: Evolution of mantle photoreceptor organs in bivalves (Bivalvia, Pteriomorphia). Evolution 74:2105–2120.

Ayalon, I., L. F. de Barros Marangoni, J. I. C. C. Benichou, D. Avisar, O. Levy, L. F. Barros Marangoni, J. I. C. C. Benichou, D. Avisar, O. Levy, L. F. de Barros Marangoni, J. I. C. C. Benichou, D. Avisar, and O. Levy. 2019. Red Sea corals under Artificial Light Pollution at Night (ALAN) undergo oxidative stress and photosynthetic impairment. Global Change Biology 25:4194–4207.

Bedrosian, T. A., C. A. Vaughn, A. Galan, G. Daye, Z. M. Weil, and R. J. Nelson. 2013. Nocturnal light exposure impairs affective responses in a wavelength-dependent manner. Journal of Neuroscience 33:13081–13087.

Bertolini, C., S. Rubinetti, G. Umgiesser, R. Witbaard, T. J. Bouma, A. Rubino, and R. Pastres. 2021. How to cope in heterogeneous coastal environments: Spatio-temporally endogenous circadian rhythm of valve gaping by mussels. Science of the Total Environment 768:145085.

Bolton, D., M. Mayer-Pinto, G. F. Clark, K. A. Dafforn, W. A. Brassil, A. Becker, and E. L. Johnston. 2017. Coastal urban lighting has ecological consequences for multiple trophic levels under the sea. Science of the Total Environment 576:1–9.

Brooks, M. E., K. Kristensen, K. J. van Benthem, A. Magnusson, C. W. Berg, A. Nielsen, H. J. Skaug, M. Mächler, and B. M. Bolker. 2017. glmmTMB Balances Speed and Flexibility Among Packages for Zero-inflated Generalized Linear Mixed Modeling. R Journal 9:378–400.

Brüning, A., W. Kloas, T. Preuer, and F. Hölker. 2018. Influence of artificially induced light pollution on the hormone system of two common fish species, perch and roach, in a rural habitat. Conservation Physiology 6.

Cho, J., J. H. Park, J. K. Kim, and E. F. Schubert. 2017. White light-emitting diodes: History, progress, and future. Laser and Photonics Reviews 11.

Clements, J. C., and L. A. Comeau. 2019. Use of High-Frequency Noninvasive Electromagnetic Biosensors to Detect Ocean Acidification Effects on Shellfish Behavior. Journal of Shellfish Research 38:811–818.

Comeau, L. A., J. M. F. Babarro, A. Longa, and X. A. Padin. 2018. Valve-gaping behavior of raft-cultivated mussels in the Ría de Arousa, Spain. Aquaculture Reports 9:68–73.

Cornwall, M. C., and A. L. F. Gorman. 1983. Ionic and spectral mechanisms of the off response to light in hyperpolarizing photoreceptors of the clam, Lima scabra. Cellular and Molecular Neurobiology 3:311–328.

Cronly-Dillon, J. R. 1966. Spectral Sensitivity of the Scallop Pecten maximus. Science 151:346–347.

Davies, T. W., M. Coleman, K. M. Griffith, and S. R. Jenkins. 2015. Night-time lighting alters the composition of marine epifaunal communities. Biology Letters 11.

Davies, T. W., D. McKee, J. Fishwick, S. Tidau, and T. Smyth. 2020. Biologically important artificial light at night on the seafloor. Scientific Reports 10:1–10.

Davies, T. W., and T. Smyth. 2018. Why artificial light at night should be a focus for global change research in the 21st century.

Diamantopoulou, C., E. Christoforou, D. M. Dominoni, E. Kaiserli, J. Czyzwski, N. Mirzai, and S. Spatharis. 2021. Wavelength-dependent effects of artificial light at night on phytoplankton growth and community structure:1–26.

Dominoni, D. M., W. Goymann, B. Helm, and J. Partecke. 2013. Urban-like night illumination reduces melatonin release in European blackbirds (Turdus merula): Implications of city life for biological time-keeping of songbirds. Frontiers in Zoology 10.

Duarte, C., D. Quintanilla-Ahumada, C. Anguita, P. H. Manríquez, S. Widdicombe, J. Pulgar, E. A. Silva-Rodríguez, C. Miranda, K. Manríquez, and P. A. Quijón. 2019. Artificial light pollution at night (ALAN) disrupts the distribution and circadian rhythm of a sandy beach isopod. Environmental Pollution 248:565–573.

Fernández, A., U. Grienke, A. Soler-Vila, F. Guihéneuf, D. B. Stengel, and D. Tasdemir. 2015. Seasonal and geographical variations in the biochemical composition of the blue mussel (Mytilus edulis L.) from Ireland. Food Chemistry 177:43–52.

Garci, R., J. R. Garcia-March, J. R. García-March, M. Á. Sanchís Solsona, A. M. García-Carrascosa, and R. Garci. 2008. Shell gaping behaviour of Pinna nobilis L., 1758 : circadian and circalunar rhythms revealed by in situ monitoring. Marine Biology 153:689–698.

Garratt, M. J., S. R. Jenkins, and T. W. Davies. 2019. Mapping the consequences of artificial light at night for intertidal ecosystems. Science of the Total Environment 691:760–768.

Gaston, K. J., J. Bennie, T. W. Davies, and J. Hopkins. 2013. The ecological impacts of nighttime light pollution: A mechanistic appraisal. Biological Reviews 88:912–927.

Gaston, K. J., T. W. Davies, S. L. Nedelec, and L. A. Holt. 2017. Impacts of Artificial Light at Night on Biological Timings. Annual Review of Ecology, Evolution, and Systematics 48:49–68.

Gnyubkin, V. F. 2010. The Circadian Rhythms of Valve Movements in the Mussel Mytilus galloprovincialis 36:419–428.

Gosling, E. 2003. Bivalve Molluscs. Fishing New Books, An imprint of Blackwell Science.

Hills, A., S. Pouil, D. Hua, and T. J. Mathews. 2020. Clearance rates of freshwater bivalves Corbicula fluminea and Utterbackia imbecillis in the presence and absence of light. Aquatic Ecology 54:1059–1066.

Hölker, F., J. Bolliger, T. W. Davies, S. Giavi, A. Jechow, G. Kalinkat, T. Longcore, K. Spoelstra, S. Tidau, M. E. Visser, and E. Knop. 2021. 11 Pressing Research Questions on How Light Pollution Affects Biodiversity. Frontiers in Ecology and Evolution 9:1–13.

Kent, F. E. A., K. S. Last, D. B. Harries, and W. G. Sanderson. 2017. In situ biodeposition measurements on a Modiolus modiolus (horse mussel) reef provide insights into ecosystem services. Estuarine, Coastal and Shelf Science 184:151–157.

Kobak, J., and P. Nowacki. 2007. Light-related behaviour of the zebra mussel (Dreissena polymorpha, Bivalvia). Fundamental and Applied Limnology / Archiv für Hydrobiologie 169:341–352.

Kyba, C. C. M., T. Kuester, A. S. De Miguel, K. Baugh, A. Jechow, F. Hölker, J. Bennie, C. D. Elvidge, K. J. Gaston, and L. Guanter. 2017. Artificially lit surface of Earth at night increasing in radiance and extent. Science Advances 3:1–9.

Lenth, R. V. 2018. emmeans: Estimated Marginal Means, aka Least-Squares Means. R package version 1.1.

Maggi, E., and L. Benedetti-Cecchi. 2018. Trophic compensation stabilizes marine primary producers exposed to artificial light at night. Marine Ecology Progress Series 606:1–5.

Miller, D. C., and E. Bretschneider. 2006. Method of Lighting for Protecting Sea Turtles 2.

Miner, K. A., M. Huertas, A. S. Aspbury, and C. R. Gabor. 2021. Artificial Light at Night Alters the Physiology and Behavior of Western Mosquitofish (Gambusia affinis). Frontiers in Ecology and Evolution 9:1–12.

Morton, B. 2008. The evolution of eyes in the Bivalvia: New insights. American Malacological Bulletin 26:35–45.

Naylor, E. 1999. Marine animal behaviour in relation to lunar phase. Earth, Moon and Planets 85–86:291–302.

Nielsen, M. V., and T. Strömgren. 1985. The effect of light on the shell length growth and defaecation rate of Mytilus edulis (L.). Aquaculture 47:205–211.

Okumus, I., and H. P. Stirling. 1998. Seasonal variations in the meat weight, condition index and biochemical composition of mussels (Mytilus edulis L.) in suspended culture in two Scottish sea lochs. Aquaculture 159:249–261.

Ortmann, C., and M. K. Grieshaber. 2003. Energy metabolism and valve closure behaviour in the Asian clam Corbicula fluminea:4167–4178.

Pampapathi, K. 1954. Tidal Rhythmicity of Rate of Water Propulsion in Mytilus, and Its Modifiability by Transplantation 106:353–359.

Pittendrigh, C. 1960. Circadian rhythms and the circadian organization of living systems. Cold Spring Harbor symposia on quantitative biology 25:159–184.

Poot, H., B. J. Ens, H. de Vries, M. A. H. Donners, M. R. Wernand, and J. M. Marquenie. 2008. Green light for nocturnally migrating birds. Ecology and Society 13.

Prather, C. M., S. L. Pelini, A. Laws, E. Rivest, M. Woltz, C. P. Bloch, I. Del Toro, C. K. Ho, J. Kominoski, T. A. S. Newbold, S. Parsons, and A. Joern. 2013. Invertebrates, ecosystem services and climate change. Biological Reviews 88:327–348.

Prins, T. C., A. C. Smaal, and R. F. Dame. 1998. A review of the feedbacks between bivalve grazing and ecosystem processes. Aquatic Ecology 31:349–359.

R Core Team. 2021. R: A Language and Environment for Statistical Computing. Vienna, Austria.

Riisgård, H. U., J. Lassen, and C. Kittner. 2006. Valve-gape response times in mussels (Mytilus edulis) - Effects of laboratory preceding-feeding conditions and in situ tidally induced variation in phytoplankton biomass. Journal of Shellfish Research 25:901–911.

Robson, A. A., G. R. Thomas, C. G. de leaniz, and R. P. Wilson. 2009. Valve gape and exhalant pumping in bivalves: Optimization of measurement. Aquatic Biology 6:191–200.

Robson, C. Garcia, R. P. Wilson, and L. G. Halsey. 2010a. Effect of anthropogenic feeding regimes on activity rhythms of laboratory mussels exposed to natural light:197–204.

Robson, C. Garcia De Leaniz, R. P. Wilson, and L. G. Halsey. 2010b. Behavioural adaptations of mussels to varying levels of food availability and predation risk. Journal of Molluscan Studies 76:348–353.

Von Salvini-Plawen, L. 2008. Photoreception and the polyphyletic evolution of photoreceptors (with special reference to Mollusca). American Malacological Bulletin 26:83–100.

Saurel, C., J. C. Gascoigne, M. R. Palmer, and M. J. Kaiser. 2007. In situ mussel feeding behavior in relation to multiple environmental factors: Regulation through food concentration and tidal conditions. Limnology and Oceanography 52:1919–1929.

van der Schatte Olivier, A., L. Jones, L. Le Vay, M. Christie, J. Wilson, and S. K. Malham. 2018. A global review of the ecosystem services provided by bivalve aquaculture. Reviews in Aquaculture 12:1–23.

Sharma, V. K. 2003. Adaptive Significance of Circadian Clocks. Chronobiology International 20:901–919.

Smithson, M., and J. Verkuilen. 2006. A better lemon squeezer? Maximum-likelihood regression with beta-distributed dependent variables. Psychological Methods 11:54–71.

Smyth, T. J., A. E. Wright, D. McKee, S. Tidau, R. Tamir, Z. Dubinsky, D. Iluz, and T. W. Davies. 2021. A global atlas of artificial light at night under the sea. Elementa 9:1–13.

Speiser, D. I., E. R. Loew, and S. Johnsen. 2011. Spectral sensitivity of the concave mirror eyes of scallops: Potential influences of habitat, self-screening and longitudinal chromatic aberration. Journal of Experimental Biology 214:422–431.

Spoelstra, K., R. H. A. Van Grunsven, M. Donners, P. Gienapp, M. E. Huigens, R. Slaterus, F. Berendse, M. E. Visser, and E. Veenendaal. 2015. Experimental illumination of natural habitat—an experimental set-up to assess the direct and indirect ecological consequences of artificial light of different spectral composition. Philosophical Transactions of the Royal Society B: Biological Sciences 370.

Strömgren, T. 1976. Length growth of Mytilus Edulis (Bivalvia) in relation to photoperiod, irradiance, and spectral distribution of light. Sarsia 61:31–40.

Tantanasarit, C., S. Babel, A. J. Englande, and S. Meksumpun. 2013. Influence of size and density on filtration rate modeling and nutrient uptake by green mussel (Perna viridis). Marine Pollution Bulletin 68:38–45.

Tidau, S., T. Smyth, D. McKee, J. Wiedenmann, C. D’Angelo, D. Wilcockson, A. Ellison, A. J. Grimmer, S. R. Jenkins, S. Widdicombe, A. M. Queirós, E. Talbot, A. Wright, and T. W. Davies. 2021. Marine artificial light at night: An empirical and technical guide. Methods in Ecology and Evolution 12:1588–1601.

Tran, D., M. Perrigault, P. Ciret, and L. Payton. 2020. Bivalve mollusc circadian clock genes can run at tidal frequency. Proceedings of the Royal Society B: Biological Sciences 287:1–9.

Wilkens, L. A. 2008. Primary inhibition by light: A unique property of bivalve photoreceptors. American Malacological Bulletin 26:101–109.

Williams, B. G., and C. A. Pilditch. 1997. The Entrainment of Persistent Tidal Rhythmicity in a Filter-Feeding Bivalve Using Cycles of Food Availability. Journal of Biological Rhythms 12:173–181.

Wood, S. 2011. Fast stable restricted maximum likelihood and marginal likelihood estimation of semiparametric generalized linear models Author (s): Simon N. Wood Published by : Wiley for the Royal Statistical Society Stable URL : http://www.jstor.org/stable/41057423. Journal of the Royal Statistical Society Series B (Statistical Methodology) 73:3–36.

Zaldibar, B., I. Cancio, and I. Marigómez. 2004. Circatidal variation in epithelial cell proliferation in the mussel digestive gland and stomach. Cell and Tissue Research 318:395–402.

Zapata, M. J., S. M. P. Sullivan, and S. M. Gray. 2018. Artificial Lighting at Night in Estuaries— Implications from Individuals to Ecosystems.

Zissis, G., and P. Bertoldi. 2018. Status of LED-Lighting world market in 2017. Joint Research Centre:76.

