## Supplementary Material for "The effects of artificial light at night (ALAN) on the gaping activity and feeding of mussels"

### S1 Tetraselmis sp. monoculture

The monoculture of the green microalgae *Tetraselmis* sp. was cultivated from an inoculum taken from microalgae disks of live *Tetraselmis* sp. (Florida Aqua Farms) and supplemented with F/2 media according to Guillard R.L. Robert (1975). The cultures were maintained at a constant illumination under fluorescent lights, temperature of 21°C and were renewed bi-weekly. The concentration of the dense culture was calculated by using Fast-Read 102® counting chambers.

### S2 LED Light spectra

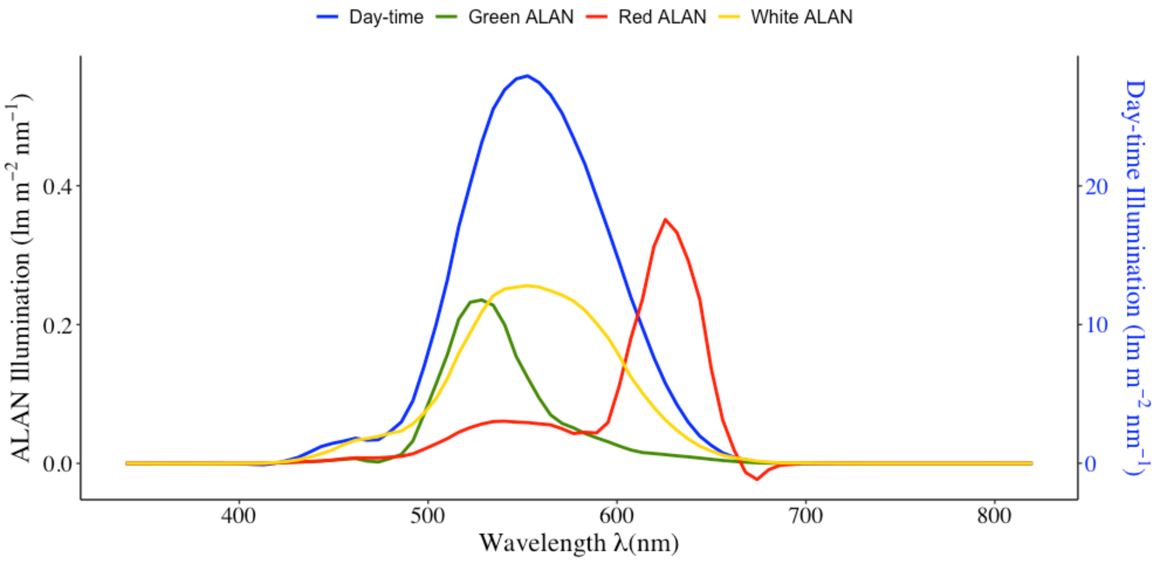


Figure S1: Spectra profile of the day (blue - right y-axis) and ALAN (green, red and white – left y-axis) measured with an Apogee® SS-110 Field Spectroradiometer. The illumination level of daylight was standardised at ~3700Lux and all ALAN treatments were standardised at ~20Lux.

### S3 Valvometry system

For each run of the experiment, 12 independent reed switches were inserted in 15ml plastic serological pipettes which were sealed to prevent water damage and connected to three 4-channel boxes. These boxes were equipped with a potentiometer which attenuated the voltage from 5 Volt-DC to 2.5 volts, as required by the data acquisition devise. For this purpose, a Picolog1216 USB Voltage Data Logger (Pico Technology®), was connected through an external terminal board, which would send the signal from the 12 reed switches. Any active reed switch supplied 2.5 volts to the Picolog devise indicating the presence of physiological activity of the mussel's shell. The Picolog6® dedicated software was used to record the signal.

Each switch could be set at close circuit position by using a small Neodynium (N42) rod magnet (Magnet Experi ®, F214, 2mm diameter x 4mm long). A magnet was attached to a thin stainless-steel strip which in turn was attached to the posterior end of the left valve of the mussel via an aquarium safe instant adhesive gel (JBL PRO HARY®). The stainless-steel strip was bent to avoid any contact with the protruding mantle at the exhalant valve. The serological pipette was secured at the right valve of the mussel via a thin strip of a rubber waterproof tape (Flex Tape®, FTB501). When the mussel was at a closed position (i.e., the magnet was at a close proximity causing the reed switch to close), 2.5 volts would be applied to one of the inputs of the PicoLog1216. When the mussel was open, there would be no output hence zero volts would be recorded. A simplified schematic representation of the mussel gape tracking system can be seen in Figure S2.

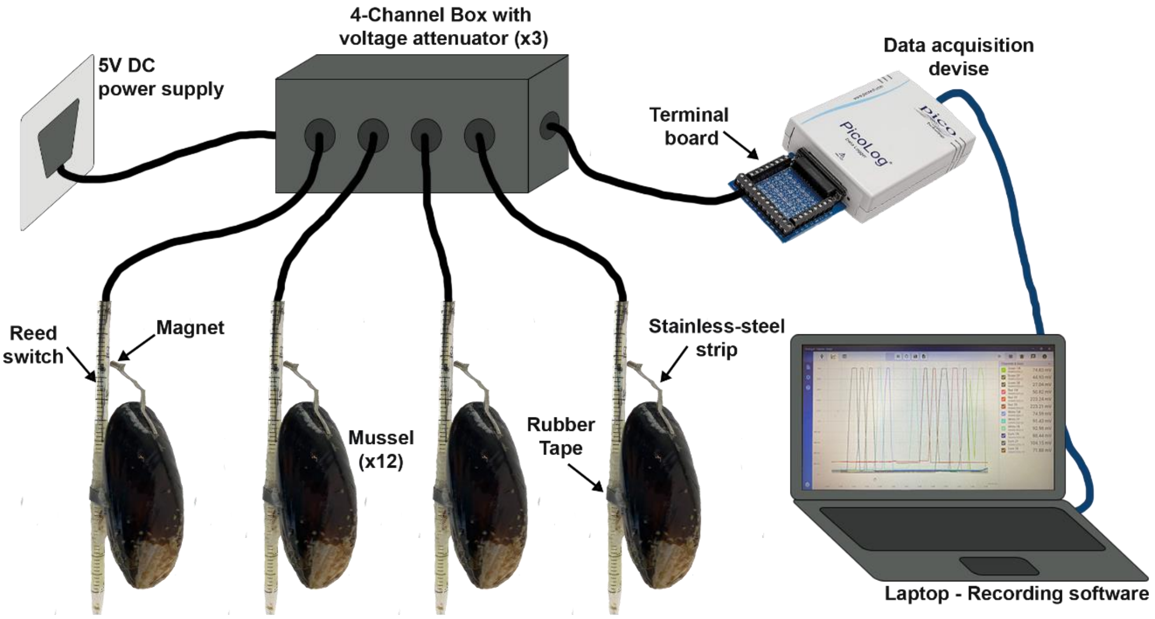


Figure S2: Simplified schematic representation of the mussel valvometry system. Only one out of the 3-in-total 4-chanel boxes are presented hence only four out of the 12 mussels tracked per run.

#### S4 Identifying and excluding outliers

The data from the phytoplankton removal capacity experiment were negatively skewed (medcouple = -0.234). Hence, an outlier was considered any value falling outside the range described by Hubert and Van Der Veeken (2008) on univariate data. Identified outliers are labelled in Figure S4 and were excluded from any analysis.

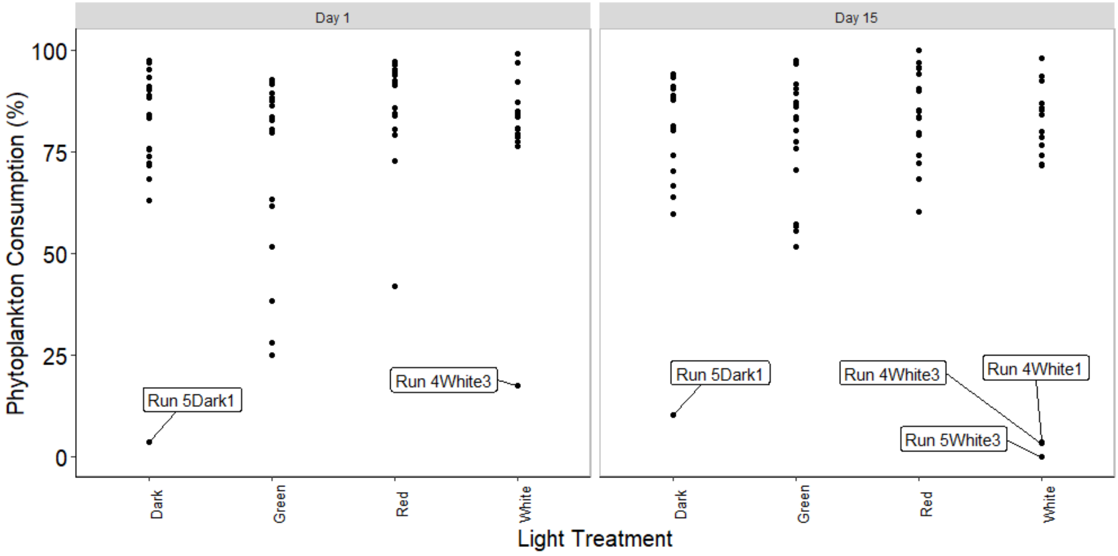


Figure S **Error! No text of specified style in document.**: The percentage of phytoplankton consumed by mussels, per experimental day, where the identified outliers are annotated with their individual mussel ID.

### S5 Model summary outputs

Table S 4-1: Detailed summary of the best supported GAMs and GLMM of the response variables (A) Proportion of time open, (B) Gaping frequency (C) Phytoplankton consumption.

**(A) Proportion of time open**

| **Explanatory Variable** | **Estimate** | **Standard Error** | **Z-Value** | **P-Value** |
|---|---|---|---|---|
| Intercept | 0.2724 | 0.0636 | 4.281 | $1.86 \times 10^{-5}$ |
| Experimental day | -0.051 | 0.003 | -16.625 | $<2 \times 10^{-16}$ |
| Water Change (Yes) | 0.300 | 0.020 | 14.772 | $<2 \times 10^{-16}$ |
| | **edf** | **Reference df** | **Chi-square** | **P-Value** |
| s(Hour):Dark | 6.212 | 8 | 76.59 | $<2 \times 10^{-16}$ |

| | | | | |
|---|---|---|---|---|
| s(Hour):Green | 5.790 | 8 | 88.89 | $<2 \times 10^{-16}$ |
| s(Hour):Red | 6.854 | 8 | 210.41 | $<2 \times 10^{-16}$ |
| s(Hour):White | 5.973 | 8 | 122.74 | $<2 \times 10^{-16}$ |
| Random effect: s(musselID, Run) | 67.857 | 70 | 2393.32 | $<2 \times 10^{-16}$ |
| Deviance explained | 28.6% | | | |
| Adjusted $R^2$ | 0.196 | | | |

### (B) Gaping Frequency

| Explanatory Variable | Estimate | Standard Error | Z-Value | P-Value |
|---|---|---|---|---|
| Intercept | -0.1701 | 0.1952 | -0.872 | 0.3833 |
| Experimental day | -0.0406 | 0.0029 | -14.229 | $<2 \times 10^{-16}$ |
| Water Change (Yes) | 0.1258 | 0.0188 | 6.694 | $2.17 \times 10^{-11}$ |
| Green (Night) | 0.3393 | 0.1372 | 2.473 | 0.0134 |
| Red (Night) | -0.3108 | 0.1570 | -1.980 | 0.0477 |
| White (Night) | -0.2433 | 0.1632 | -1.491 | 0.1359 |

| | edf | Reference df | Chi-square | P-Value |
|---|---|---|---|---|
| s(Hour):Dark | 6.971 | 8 | 108.1 | $<2 \times 10^{-16}$ |
| s(Hour):Green | 4.953 | 8 | 107.1 | $<2 \times 10^{-16}$ |
| s(Hour):Red | 6.374 | 8 | 109.9 | $<2 \times 10^{-16}$ |
| s(Hour):White | 6.906 | 8 | 144.6 | $<2 \times 10^{-16}$ |
| Random effect: s(musselID, Run) | 65.881 | 67 | 5791.2 | $<2 \times 10^{-16}$ |
| Deviance explained | 22% | | | |
| Adjusted $R^2$ | 0.162 | | | |

### (C) Phytoplankton consumption

| Explanatory Variable | Estimate | Standard Error | Z-Value | P-Value |
|---|---|---|---|---|

| | | | | |
|---|---|---|---|---|
| Intercept | 1.0280 | 0.3859 | 2.664 | 0.0077 |
| Proportion open:Green | -2.5958 | 0.9064 | -2.864 | 0.0042 |
| Proportion open:Red | 0.1671 | 0.7740 | 0.216 | 0.8291 |
| Proportion open:White | 1.2633 | 0.8195 | 1.542 | 0.1232 |

### S6 Mussel gaping activity compared with the tidal pattern

Tidal data were provided by the British Oceanographic Data Centre (BODC). As data for Loch Eil were not available, the data used were from Tobermory, Scotland, the closest location to the mussel collection site. The tidal height at the dates the experiments were running were compared with the mussel activity, both averaged in a 24hour period (Figure S7).

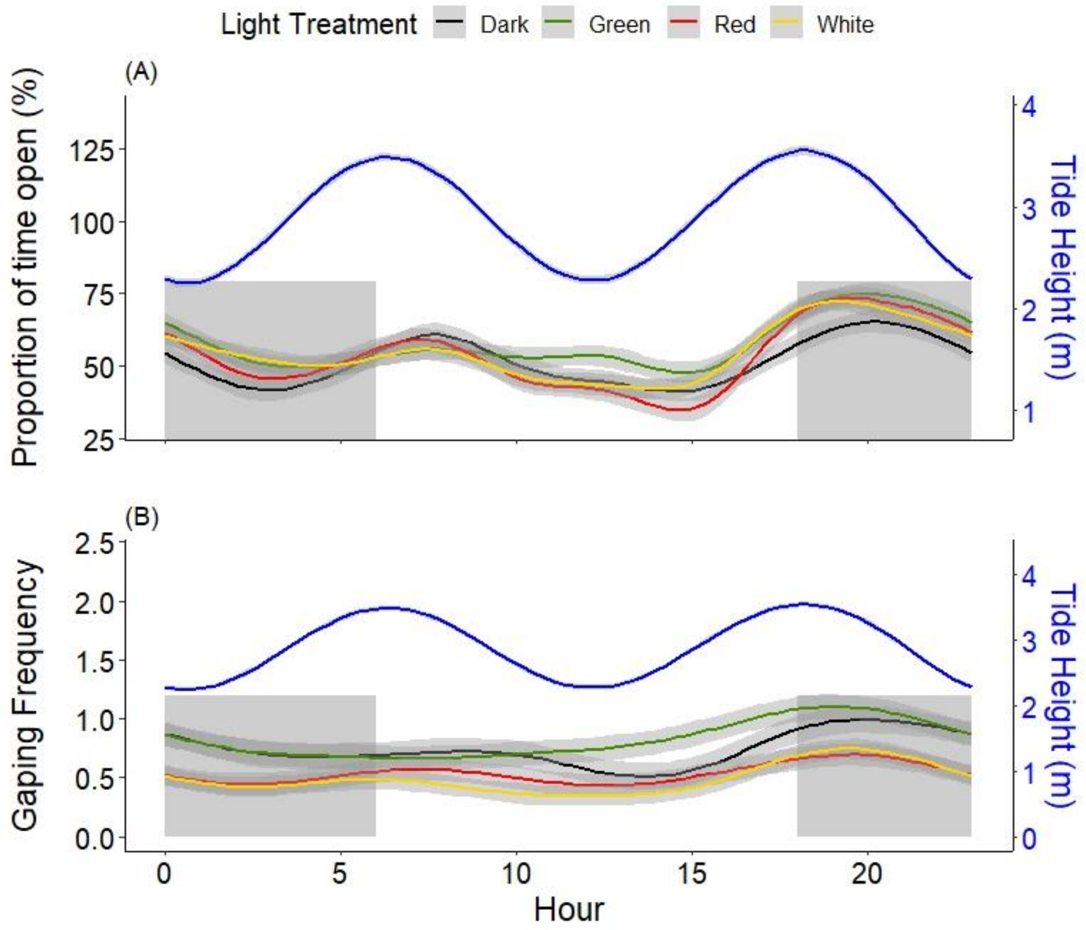


Figure S7**:** The average (lines) and standard error (shading of lines) of the (A) proportion of open gape expressed as a percentage and (B) open/close frequency at each light treatment (dark, green, red, and white) through a 24hour period. The white and grey sections represent day-time and night-time respectively. The blue line (right y-axes) plots the hourly tidal height (m).
